# SUMO E1 covalent allosteric inhibitors modulate polyamine synthesis via the MAT2A-AdoMetDC axis

**DOI:** 10.1101/2024.12.12.627095

**Authors:** Shuai Zhang, Zhiying Wang, Yutong Jiao, Xiaoyu Shi, Yuanbao Ai, Juanping Wang, Sen Liu

## Abstract

Upregulation of protein SUMOylation is associated with various diseases, and SUMOylation inhibitors are promising drug candidates. We performed the first virtual screening of SUMO E1 covalent allosteric inhibitors (CAIs) and identified two SUMO E1 CAIs with new scaffolds and covalent warheads. We further demonstrated that these new CAIs perturbed the SUMOylation pathway and protein SUMOylation. Specifically, these CAIs affected the SUMOylation of the methionine adenosyltransferases MAT2A. The inhibition of MAT2A SUMOylation unexpectedly stimulated polyamine synthesis. Lastly, we showed that the combination of SUMO E1 CAIs with a polyamine synthesis inhibitor had synergistic effects in inhibiting T47D cells. Our work demonstrated the first cost-effective virtual screening of SUMO E1 CAIs, found that the downregulation of MAT2A SUMOylation increases polyamine synthesis, and SUMO E1 CAIs can synergize with polyamine synthesis inhibitors. This work would be of great value to the study of SUMOylation, covalent/allosteric drugs, and the polyamine metabolism network.

## Introduction

Post-translational modifications modulate various protein functions under both normal and disease states ^1,2^. One of the well-known protein post-translational modifications is ubiquitylation, which attaches ubiquitin to target proteins for proteasome dependent degradation ^3^. SUMOylation is a ubiquitin-like (Ubl) post-translational process attaching the SUMO (small ubiquitin-like modifier) protein to target proteins ^4^. SUMOylation also modulates various cellular functions, such as signal transduction, gene transcription, oncogenesis, and immune response ^4,5^. Closely resembling the ubiquitylation pathway, the SUMOylation pathway is a process catalyzed by an enzymatic cascade consisting of three classes of enzymes (Figure 1a): SUMO E1, SUMO E2, and SUMO E3 ^4^. SUMO E1 is an ATP-dependent SUMO-activating enzyme (SAE), E2 is a SUMO conjugating enzyme, and E3 is a SUMO ligase. A dramatic difference between the ubiquitylation pathway and the SUMOylation pathway is that only one E2 enzyme (Ubc9) has been identified in humans thus far ^4^. In addition, contrary to ubiquitylation’s function in inducing protein degradation, SUMOylation usually stabilizes target proteins and enhances protein-protein interactions ^4^.

**Figure 1.**
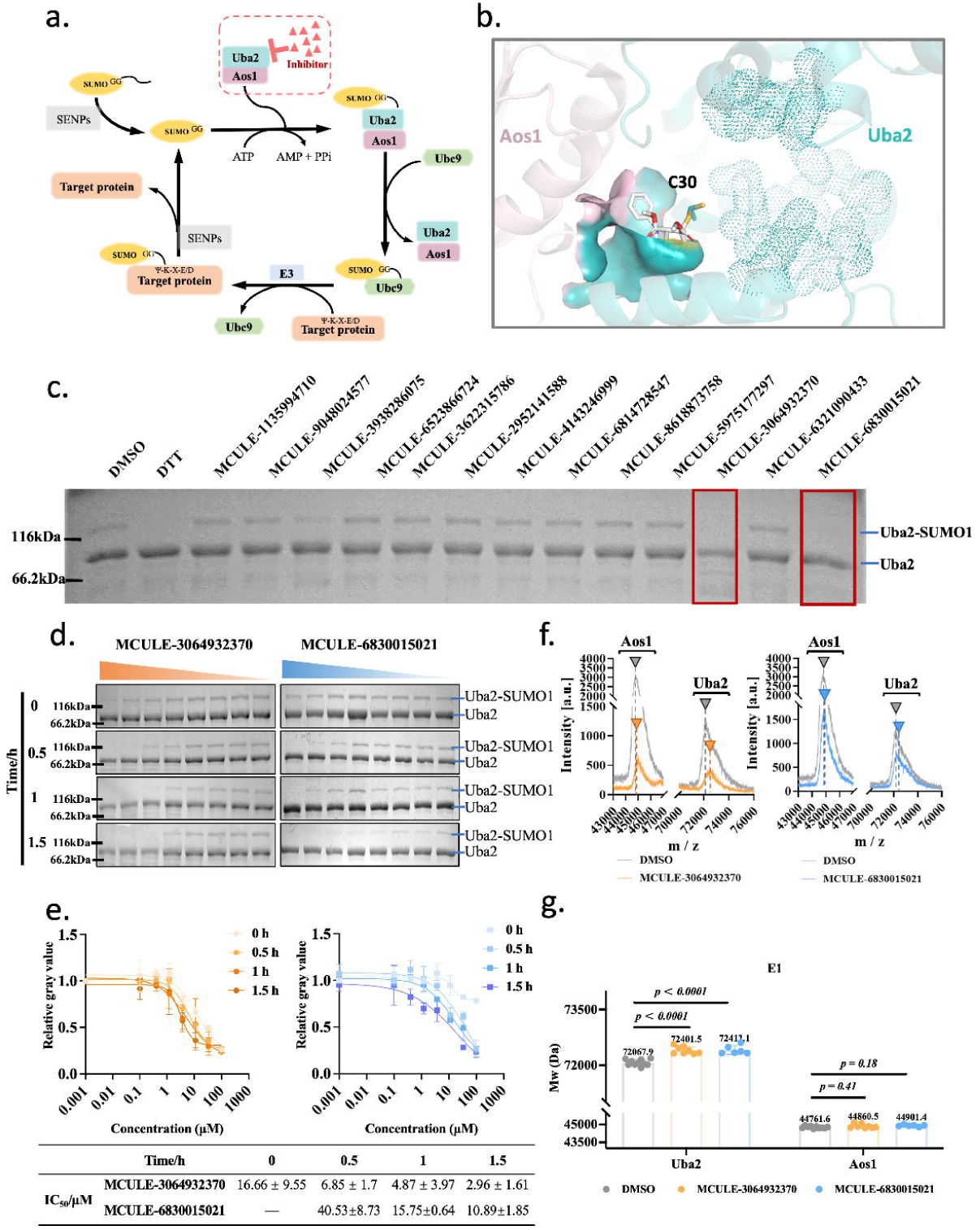
Screening of new covalent inhibitors targeting the allosteric pocket of SUMO E1. (a) A schematic diagram of the SUMOylation pathway. SUMO E1 is a heterodimer of Aos1 and Uba2. Ubc9 is the sole SUMO E2 discovered thus far. (b) The complex of SUMO E1 and a covalent allosteric inhibitor (CAI) COH000 (PDB ID: 6CWY). COH000 is shown in sticks and formed a covalent bond with Uba2-C30. The residues of the allosteric pocket are shown in surface, and the residues of the ATP-binding pocket are shown in dotted surface. The SUMO-binding site is far away from the allosteric pocket and not shown. (c) MCULE-3064932370 and MCULE-6830015021 inhibited the formation of the SUMOylation of Uba2. Before the addition of SUMO1, the compounds were pre-incubated with SUMO E1 (Aos1/Uba2) at 37 °C for 30 minutes at various concentrations according to their solubility. MCULE-3064932370 and MCULE-6830015021 were 100 µM, MCULE-6321090433 was 50 µM, and the others were 500 µM. The reductant DTT (dithiothreitol) inhibits SUMOylation and was used as a control. The results with 60 min and 90 min pre-incubation had similar results (Figure S1, S2). All samples had 1% (v/v) DMSO. (d) Time-dependent and concentration-dependent inhibition of MCULE-3064932370 and MCULE-6830015021 on the SUMOylation of Uba2. The quantitative data from three replicates are shown in (e). Error bars represent mean ± s.d.. (f) Representative spectra from the full-protein mass spectrometry analysis. (g) Statistical analysis of the m/z data from multiple MALDI-TOF detections (n ≥ 6).

The dysregulation of the SUMOylation pathway leads to the dysregulation of the SUMOylation of downstream protein targets, among which are c-Myc^6^, Tau^7^, and NOTCH^8^. In total, it has been discovered that around 200-4000 proteins in human cell might be regulated by SUMOylation ^9,10^. The SUMOylation pathway is dysregulated in various diseases including cancer, neurodegeneration, and infection ^4^. Therefore, the SUMOylation pathway has attracted increasing attentions in drug discovery, and inhibitors targeting SUMO E1 such as natural products and synthetic compounds have been reported ^11–18^. SUMO E1 is a heterodimer formed by Aos1 (or SAE1) and Uba2 (or SAE2), and the catalytic pocket mostly locates in the Uba2 subunit (Figure 1b). ML-792 and TAK-981 are extensively optimized mechanism-based inhibitors binding to the ATP-binding site of the catalytic pocket and forming covalent SUMO-inhibitor adducts with a sulfamate warhead (covalent group) ^17,18^. The clinical trials of TAK-981 (subasumstat) have shown promising results in treating various cancers ^5^. In another scenario, COH000 was discovered by experimentally screening 290,921 compounds and additional chemical optimization ^11,19^. Different from ML-792 and TAK-981, COH000 covalently attached to Uba2-C30 located in an allosteric pocket (Figure 1b) via a 7-oxabicyclo[2.2.1]hept-2-ene warhead. This allosteric pocket is concaved by the two subunits (Aos1/Uba2), away from the ATP binding site and the catalytic pocket ^19^, so the inhibition of COH000 is not competed by ATP or SUMO ^11^. COH000 also showed high specificity, since this allosteric pocket depends on the conformational changes of Aos1/Uba2 and was not noticed in other Ubl E1 enzymes ^11^. Therefore, discovering more allosteric covalent SUMO E1 inhibitors would be of great value.

The development of the covalent SUMO E1 inhibitors ML-792, TAK-981, and COH000, has been echoed with the resurgence of covalent drugs in the last decade ^20^. Covalent drugs were intentionally avoided in traditional drug development pipelines due to potential idiosyncratic toxicity ^21^. Nonetheless, a retrospective analysis revealed the surprising benefits of covalent drugs ^22^, and recent years have witnessed the enormous value of targeted covalent inhibitors (TCIs) from the approvement of several blockbuster covalent drugs including sotorasib targeting K-Ras-G12C, ibrutinib targeting BTK, osimertinib targeting EGFR-T790M, and nirmatrelvir targeting the SARS-CoV2 main protease ^20^. To facilitate the discovery of covalent drugs, our lab presented SCARdock^23–27^, a computational high-throughput covalent drug screening pipeline based on the theory that the pre-requisite step for TCIs is the non-covalent binding step, which positions the warhead for covalent bond formation. SCARdock was the first computational screening tool for TCIs based on non-covalent docking ^28–31^. It was also one of the first computational covalent drug screening protocols successfully validated by experimental work ^27^. Encouraged by the fast growth of covalent drugs, many computational screening tools have been developed in recent years ^29–31^. However, only a handful tools have been experimentally validated, and more importantly, none has been confirmed to be applicable in discovering covalent drugs targeting protein-protein interaction interfaces as well as allosteric pockets.

In this work, we set out to discovery new covalent allosteric inhibitors (CAIs) by expanding the application of SCARdock in discovering covalent inhibitors targeting protein-protein interaction interfaces and allosteric pockets. SUMO E1 was chosen because it is a heterodimeric enzyme with an allosteric pocket locating at the PPI interface of two subunits (Figure 1b). We successfully identified two CAIs of SUMO E1 with scaffolds and warheads different from COH000. We verified that these CAIs bind to the allosteric pocket, and their inhibition depends on the interaction between Aos1 and Uba2. We further showed that these CAIs can perturb the SUMOylation pathway in cells. At last, we demonstrated that these CAIs stimulated polyamine synthesis through the MAT2A-AdoMetDC axis and thus were synergistic with AdoMetDC inhibitors in inhibiting cancer cells.

## Results

### 1. Screening of new covalent inhibitors targeting the allosteric pocket of SUMO E1

The computational high-throughput screening was based on the complex structure of COH000 and the human SUMO E1 (PDB ID: 6CWY ^19^). For simplicity, Aos1/Uba2 and SUMO E1 is exchangeable in this paper. The concaved pocket by the two subunits was used (Figure 1b), and the covalent residue Cys30 was virtually mutated to glycine in the SCARdock protocol as previously described ^23,27,28,32^. We prepared a pre-filtered dataset containing 106,952 compounds from the MCULE screening library (https://mcule.com) based on manually curated warheads ^23^ before it was used for docking evaluation. Based on the lowest docking score, conformation ranking, covalent atom distance, and score density ^27,28^, 2,026 virtual hits were obtained for visual verification. After discarding hits with misaligned warheads and considering warhead diversity, we chose 25 hits for purchase inquiry from commercial vendors. Finally, we were able to purchase 13 compounds with eight different warheads for experimental validation (Table S1).

An SDS-PAGE based in vitro assay ^11,19^ was used to evaluate if these compounds inhibit SUMO E1’s activity. Without inhibitors, the mixing of SUMO E1 and SUMO1 leads to the SUMOylation of Uba2 (Uba2-SUMO1), resulting in a protein band above the Uba2 band. With an inhibitor existing, the Uba2-SUMO1 band fades or disappears. When pre-incubated with SUMO E1 for 30-90 minutes at 37 °C, compounds MCULE-3064932370 and MCULE-6830015021 significantly inhibited the formation of Uba2-SUMO1 (Figure 1c, Figure S1, S2). To confirm if these two compounds are covalent inhibitors, we did time-dependent and concentration-dependent experiments. The data showed that the ratio of Uba2-SUMO1 was concentration dependent, and the IC_50_ values decreased when the incubation time increased (Figure 1d & 1e). Furthermore, we used full-protein mass spectrometry to check if these compounds covalently bind to Uba2. The MALDI-TOF data showed that both compounds covalently bound with Uba2, leading to the increase of the molecular weight of Uba2 (Figure 1f & 1g). As a reference, the molecular weight of Aos1 had no changes (Figure 1f & 1g).

### 2. The new inhibitors bind to the allosteric pocket of SUMO E1

To confirm that the new inhibitors bind to the allosteric pocket and form covalent bonds with the residue Cys30 of Uba2, we prepared the mutant protein Uba2-C30S. Similar to the previously report ^11^, we noticed that Aos1/Uba2-C30S had low activity (Figure 2a). In the time-dependent experiment (Figure 2a), both compounds showed insignificant time-dependent inhibition on the activity of Aos1/Uba2-C30S, as evaluated by the SUMOylation of Uba2-C30S (the Uba2-C30S-SUMO conjugate). In the concentration-dependent experiment (Figure 2a), MCULE-3064932370 showed inhibition at over 1.0 µM, and MCULE-6830015021 showed limited inhibition up to 1.0 mM. Considering that Ser (S) is also a residue with the potency to form covalent bonds with inhibitors, we made another Uba2 mutant, Uba2-C30A. We noticed that this mutant still had weak activity, however, the time-dependent and concentration-dependent experiments showed that the new inhibitors had far less inhibition on the SUMOylation of Uba2-C30A than the wild-type Uba2 and the Uba2-C30S mutant (Figure 2b). One possibility for this difference is that Ser (S) is more like Cys (C) than Ala (A), so the inhibitors could form better interactions. Another possibility is that the inhibitors might covalently bind to Uba2-C30S. To verify that possibility, we did full-protein mass spectrometry with these two mutants and did not notice significant changes in the molecular weights of Aos1 or Uba2 (Figure 2c, 2d).

**Figure 2.**
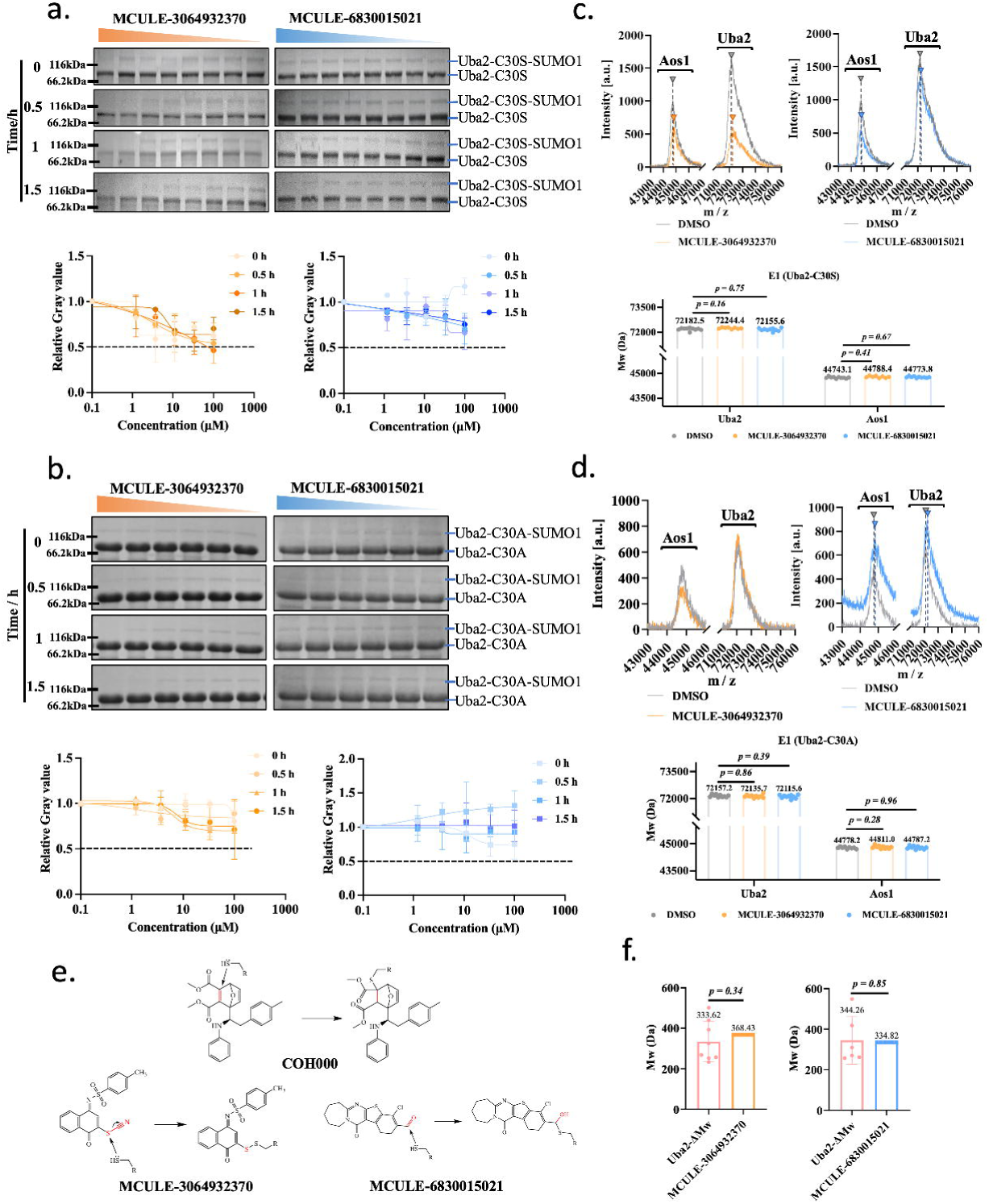
The new inhibitors bind in the allosteric pocket of SUMO E1. (a) Time-dependent and concentration-dependent inhibition of MCULE-3064932370 and MCULE-6830015021 on the SUMOylation of Uba2-C30S. The quantitative data from three replicates are shown below the representative SDS-PAGE gels. Error bars represent mean ± s.d.. (b) Time-dependent and concentration-dependent inhibition of MCULE-3064932370 and MCULE-6830015021 on the SUMOylation of Uba2-C30A. The quantitative data from three replicates are shown below the representative SDS-PAGE gels. Error bars represent mean ± s.d.. (c) Representative spectra from the full-protein mass spectrometry analysis of the Aos1/Uba2-C30S enzyme. Statistical analysis data from multiple MALDI-TOF detections (n ≥ 8) are shown below the representative spectra. (d) Representative spectra from the full-protein mass spectrometry analysis of the Aos1/Uba2-C30A enzyme. Statistical analysis data from multiple MALDI-TOF detections (n ≥ 9) are shown below the representative spectra. (e) The molecular structures and possible bonding mechanisms of MCULE-3064932370 and MCULE-6830015021 to Uba2. COH000 is shown for comparison. (f) The molecular weight differences from the full-protein spectrometry data as shown in Figure 1f & 1g.

Taken together, our data confirmed that we had discovered two new covalent allosteric inhibitors (CAIs) of the SUMO E1 enzyme, which were the first ones of this kind with the help from high-throughput virtual screening. More importantly, these two new inhibitors have non-covalent scaffolds and covalent warheads very different from the known one, i.e., COH000 (Figure 2e). MCULE-3064932370 has a thiocyanate warhead and MCULE-6830015021 has an aldehyde warhead. These warheads were widely utilized in previous covalent inhibitors ^33–35^. In accordance with the corresponding reaction mechanisms, the mass spectrometry data showed molecular weight differences close to the theoretical data (Figure 2f), confirming the covalent binding of these two ligands to Uba2.

### 3. The binding of the new SUMO E1 CAIs depends on the Aos1/Uba2 complex

Since the allosteric pocket of SUMO E1 is concaved by Aos1 and Uba2 (Figure 1b), we asked how the binding of the CAIs depends on and affects the heterocomplex. Firstly, we incubated MCULE-3064932370 and MCULE-6830015021 with Uba2 or Aos1 alone and did not notice any covalent binding of the inhibitors with either protein (Figure 3a, 3b), indicating that the binding of these inhibitors depends on the binary interaction of Aos1/Uba2. This data also hinted that the covalent binding of these two CAIs with Uba2 is not promiscuous. Then we checked how the binding of these CAIs affects the stability of the Aos1/Uba2 complex. The stability of the SUMO E1 complex was assessed with a gel-based thermal shift assay. The Aos1/Uba2 complex was pre-incubated with the CAIs before being incubated at various temperatures, and the data from quantifying the separated proteins on SDS-PAGE gels (Figure 3c) showed that both CAIs improved the thermal stability of Uba2, whereas MCULE-3064932370 also improved the thermal stability of Aos1. Next, we tested if these CAIs compete with the binding of SUMO1. Similar to COH000, the new inhibitors did not compete with SUMO1 and the concentration increase of the latter could not antagonize their inhibiting effects (Figure 3d), which is an attracting advantage of allosteric inhibitors. We noticed that the difference between MCULE-3064932370 and MCULE-6830015021 on the thermal stabilization of the Aos1/Uba2 complex could be explained from their interaction with the proteins as shown in the modeled complex structures (Figure 3e). MCULE-3064932370 forms more interactions with the residues from both Aos1 and Uba2 than MCULE-6830015021. Encouragingly, both CAIs bind to the Aos1/Uba2 complex with similar residues as COH000 does.

**Figure 3.**
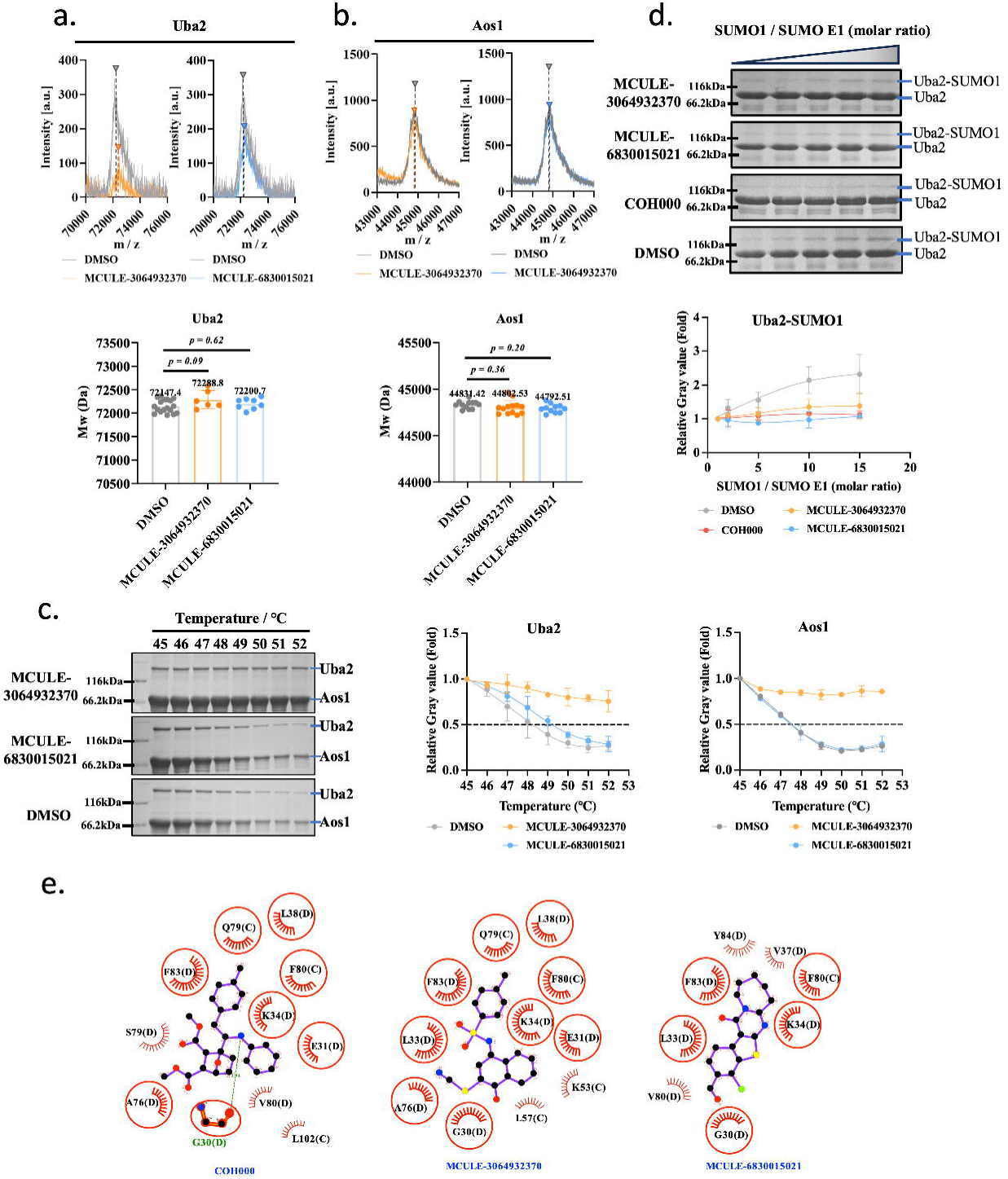
The binding of the new SUMO E1 CAIs depends on the Aos1/Uba2 complex. (a) Representative spectra and statistical analysis data from multiple MALDI-TOF detections (n ≥ 6) of the full-protein mass spectrometry analysis of the Uba2 protein when it was incubated with the inhibitors alone. (b) Representative spectra and statistical analysis data from multiple MALDI-TOF detections (n ≥ 6) of the full-protein mass spectrometry analysis of the Aos1 protein when it was incubated with the inhibitors alone. (c) The thermal denaturation result from the gel-based thermal shift assay of the Aos1/Uba2 complex with the new CAIs. (d) The competition experiment of the CAIs with the substrate SUMO1. (e) The 2D interaction diagrams of Aos1/Uba2 with COH000 (PDB ID: 6CWY), MCULE-3064932370, and MCULE-6830015021. Aos1 is the chain C, and Uba2 is the chain D. Since SCARdock only considers non-covalent interaction, the Uba2-C30 was computationally mutated to G30 as shown in the diagrams. Therefore, the ligands are shown not covalently bonded to Uba2. The ligands and residues are shown in ball-and-stick representation. The green dotted line indicates a hydrogen bond. The spoked arcs represent residues making nonbonded contacts with the ligands. The red circles indicate residues in equivalent 3D positions when the structural models are superposed.

### 4. The new SUMO E1 CAIs perturb the SUMOylation pathway

Next, we asked if these two CAIs could perturb the SUMOylation pathway in cells. Usually, SUMOylation is highly dynamic, and a very small fraction of a target protein is SUMOylated, so endogenous SUMOylated proteins are hard to be directly detected ^17^. An exception is RanGAP1, which is a well-known SUMOylation target with a high level of SUMOylation ^4^. We detected the cellular level of SUMOylated RanGAP1 in T47D breast cancer cells, which decreased upon inhibitor treatment (Figure 4a, Figure S3, S4). Accordingly, an increase of the free SUMO1 protein was found (Figure 4a, Figure S5), although we were not able to detect the SUMOylated Uba2 and Ubc9 (Figure 4b, Figure S6, S7, S8). However, we did notice that the cellular levels of Uba2 and Ubc9 decreased upon inhibitor treatment. An explanation is that SUMOylation stabilizes target proteins, so inhibition of SUMO E1 leads to the accelerated degradation of Uba2 and Ubc9.

**Figure 4.**
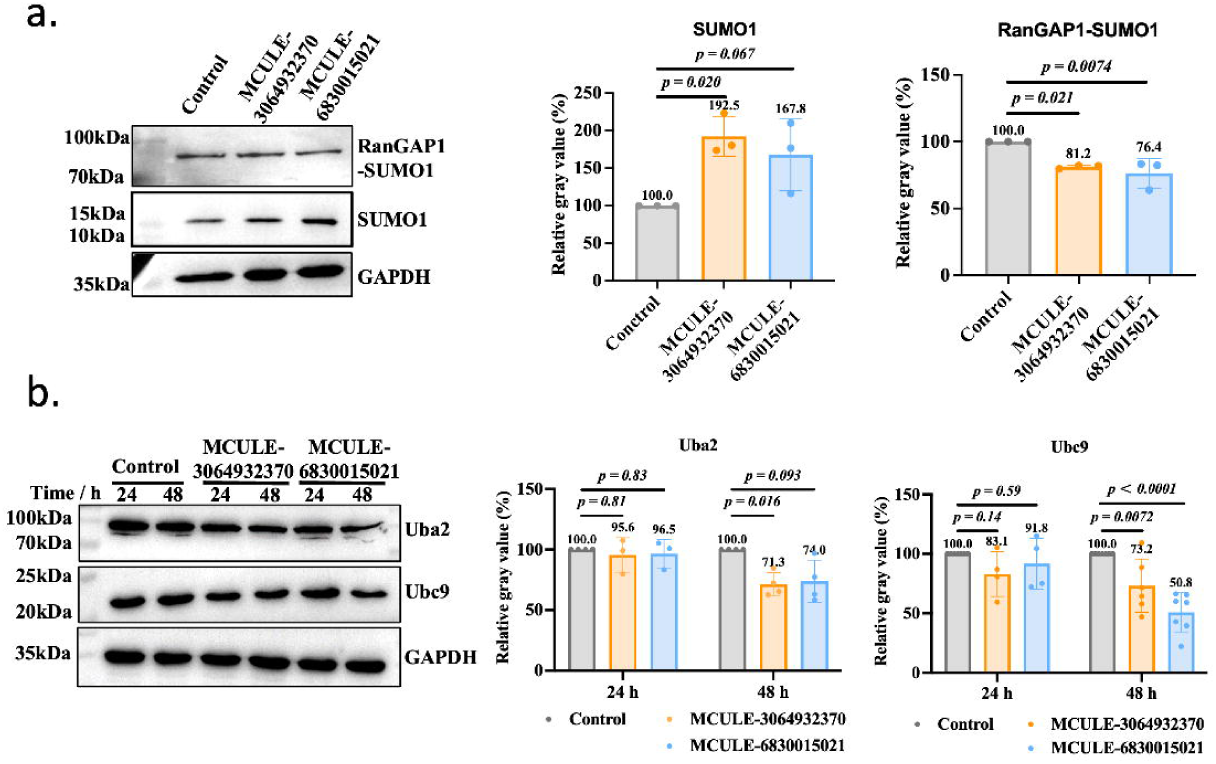
The new SUMO E1 CAIs perturb the SUMOylation pathway. T47D cells were treated with MCULE-3064932370 and MCULE-6830015021 for 48 h before evaluation. (a) The cellular levels of RanGAP1, RanGAP1-SUMO, and SUMO1 were assessed by Western blotting. (b) The cellular levels of Uba2 and Ubc9 were assessed by Western blotting.

### 5. The new SUMO E1 CAIs perturb polyamine synthesis through the MAT2A-AdoMetDC axis

Methionine adenosyltransferases (MATs) are enzymes synthesizing S-adenosylmethionine (AdoMet or SAMe), an essential player in various cellular functions and the main methyl provider in cell for the methylation of nucleic acids, proteins, lipids, and metabolites. MAT2A is wildly distributed in various kinds of cells and acts as a rate-limiting enzyme in the methionine cycle ^36^. MAT2A is also a promising therapeutic target in various cancers ^37^. As a target of the SUMO E1 enzyme, the SUMOylation of MAT2A is critical for its stability and function ^36^ (Figure 5a). To confirm that MAT2A was affected by our new CAIs, we checked its expression with Western blotting and found that MAT2A was downregulated upon the treatment of these two inhibitors (Figure 5b, Figure S9). Furthermore, the cellular level of its product AdoMet also decreased significantly (Figure 5c).

**Figure 5.**
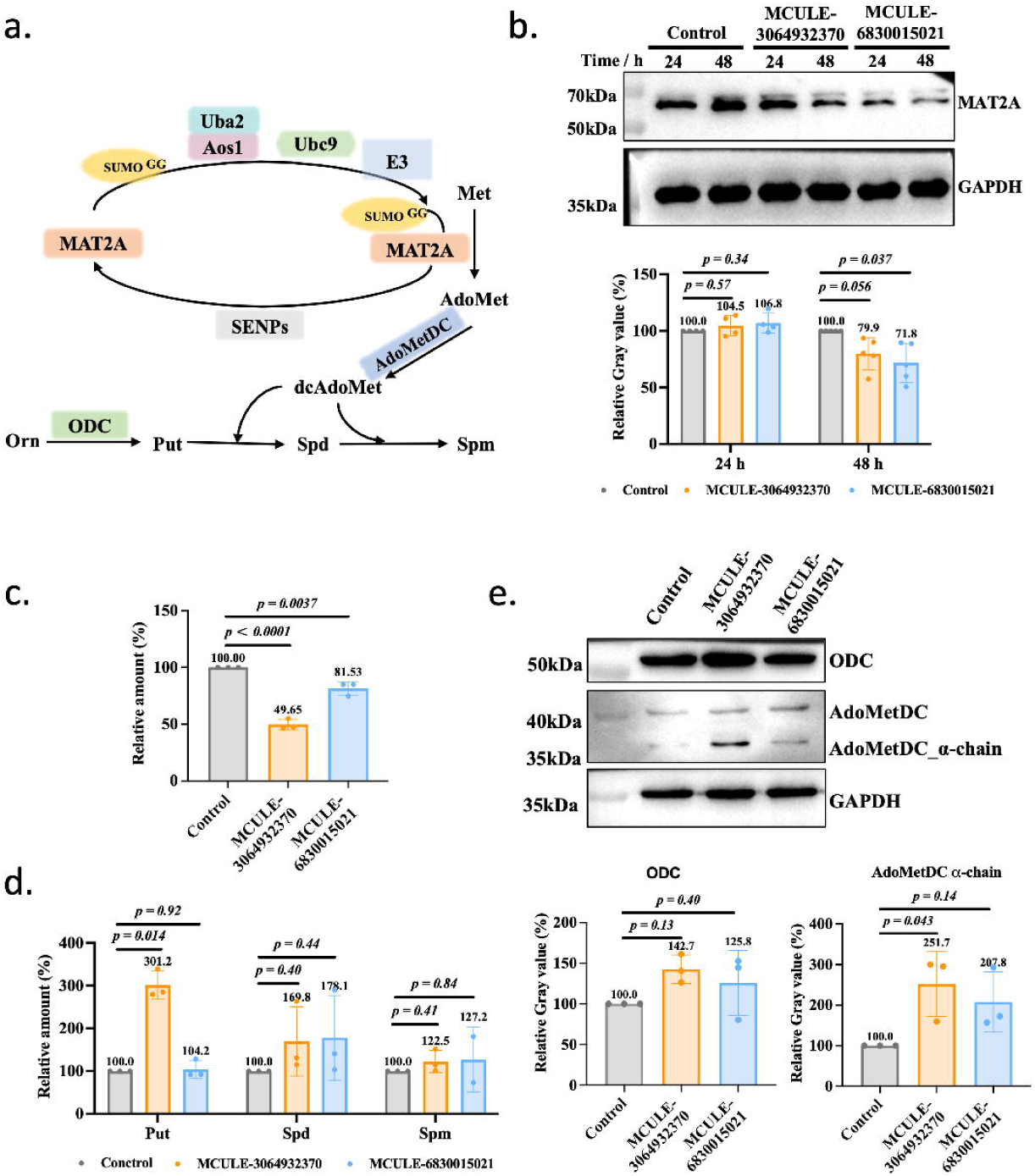
The new SUMO E1 CAIs perturb polyamine synthesis through the MAT2A-AdoMetDC axis. (a) MAT2A is a SUMOylation substrate and converts methionine (Met) to S-adenosylmethionine (AdoMet). AdoMetDC decarboxylates AdoMet to produce dcAdoMet for the synthesis of spermidine (Spd) and spermine (Spm). ODC decarboxylates ornithine (Orn) to generate putrescine (Put). (b) The cellular level of MAT2A was assessed with Western blotting after T47D cells were treated with the CAIs. (c) The cellular level of AdoMet was assessed with LC-MS after T47D cells were treated with the CAIs. (d) The cellular levels of polyamines were assessed with LC-MS after T47D cells were treated with the CAIs. (e) The cellular levels of two rate-limiting polyamine synthesis enzymes, ODC and AdoMetDC, were assessed with Western blotting after T47D cells were treated with the CAIs.

A downstream utilization of AdoMet is for producing decarboxylated AdoMet (dcAdoMet), which is catalyzed by AdoMet decarboxylase (AdoMetDC) and used for the synthesis of spermidine and spermine, two organic polyamines critical for normal cell function and increased significantly in cancer cells ^26,27,38^. To see if the decrease of AdoMet leads to the decline of cellular polyamines, we quantified the polyamine content in the breast cancer cell line T47D. Surprisingly, upon inhibitor treatment, the cellular polyamines were increased (Figure 5d). A hypothesis was that the cells compensatively upregulated polyamine synthesis. Therefore, we checked the expression levels of two rate-limiting enzymes for polyamine synthesis, ODC (ornithine decarboxylase) ^39^ and AdoMetDC. In agreement with our hypothesis, the result (Figure 5e, Figure S10, S11, S12) showed that these two enzymes were moderately upregulated upon treatment. Furthermore, the fold changes of AdoMetDC expression level and spermidine were similar (1.6x). The fold changes of spermine were lower (1.2x) but similar between both compounds.

### 6. The new SUMO E1 CAIs synergize with the AdoMetDC inhibitor in inhibiting cancer cells

Our results above showed that the inhibition of the SUMOylation pathway downregulated the cellular level of MAT2A. We analyzed the survival data of 2,277 patients with Luminal A breast cancer and found that high MAT2A level was beneficial and associated with high recurrence-free survival probability (Figure 6a). The decrease of MAT2A upregulated cellular polyamine level, but elevated polyamine level benefits cancer cells. Therefore, we reasoned that the compensative upregulation of cellular polyamine level might play a negative role upon the treatment of cancer cells by SUMO E1 inhibitors. Further dataset analysis showed that breast cancer cells have the highest dependency on MAT2A (Figure 6b), so we chose T47D (Luminal A) breast cancer cell line to assess if the new SUMO E1 CAIs synergize with polyamine synthesis inhibitors in inhibiting cancer cell growth. We found that both MCULE-3064932370 and MCULE-6830015021 inhibited the growth and proliferation of T47D cells (Figure 6c). They also inhibited the migration of T47D cells (Figure 6d, Figure S13). MCULE-3064932370 showed synergy with the AdoMetDC inhibitor MGBG (Figure 6e). Although MCULE-6830015021 did not show obvious synergy with MGBG per Chou’s CI definition ^40^, the combination treatment indeed caused higher inhibition at high concentrations (Figure 6f), indicating improved efficacy and some synergy ^41^. To gain a further understanding of the difference of these two SUMO E1 inhibitors, we performed a comparative proteomics study. The proteomics data (Figure 6g, Table S2) showed that these two inhibitors indeed caused different changes. Briefly, eight proteins were uniquely changed (upregulated) with the treatment of MCULE-3064932370, and fourteen proteins were uniquely changed (11 upregulated, 3 downregulated) with the treatment of MCULE-6830015021. The protein downregulated by both inhibitors was CEP44 (Centrosomal protein 44), which was identified as a SUMOylation target previously ^42,43^. Besides, AKR1C3 (upregulated by MCULE-3064932370) ^42,44^, ZNF384 (upregulated by MCULE-6830015021) ^42,43^, and TBC1D31 (downregulated by MCULE-6830015021) ^42,44^ were also identified as SUMOylation targets previously. In addition, another seventeen proteins (TNFRSF10A, ADCY5, OAS1, KRT1, ISG15, CYP1A2, CCK, IFIT2, MX1, OAS2, IRF9, CYP1B1, CYHR1, SPPL2A, RNF138, TMEM45B, TXN2) are also predicted SUMOylation targets (Table S3) ^45^. The detailed mechanism of the differences of these two new inhibitors would need further investigations.

**Figure 6.**
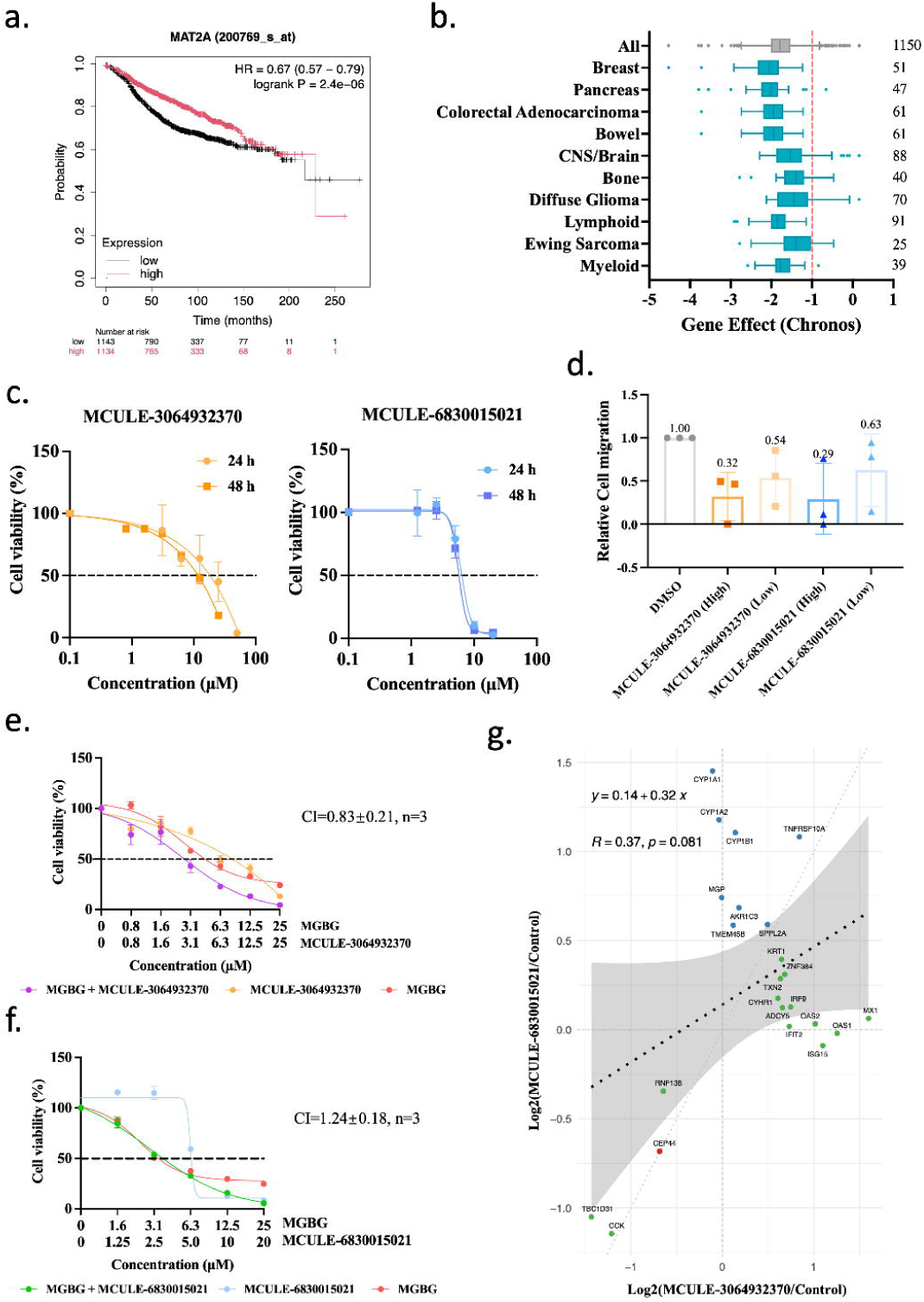
The new SUMO E1 CAIs synergize with the AdoMetDC inhibitor in inhibiting cancer cells. (a) High MAT2A expression was associated with low recurrence-free survival probability in Luminal A breast cancer patients. The data were obtained from DepMap (https://depmap.org/portal/). (b) The MAT2A CRISPR Dependence Scores (CRISPR (DepMap Public 24Q2+Score, Chronos)) of the top ten cancer types were retrieved from the DepMap ^49^. The red dotted line (−1) indicates the median Gene Effect (Chronos) −1. The case numbers are shown on the right. (c) The new CAIs inhibited the proliferation and growth of T47D cells. (d) The new CAIs inhibited the migration of T47D cells. For MCULE-3064932370: high, 25 μM; low: 12.5 μM. For MCULE-6830015021: high, 10 μM; low: 5 μM. (e) The new CAIs synergized with MGBG in inhibiting the proliferation and growth of T47D cells. (f) The comparison of the proteomic changes when T47D cells were treated by MCULE-3064932370 (20 μM) and MCULE-6830015021 (10 μM). Blue dots: upregulated by MCULE-3064932370; green dots: upregulated by MCULE-6830015021; red dot: downregulated by both.

## Discussion

SUMOylation is one of the most important protein post-translational modulation events. Thus far, it was estimated that over 4,000 proteins might have SUMOylation ^9,10^. SUMOylation regulates diverse protein functions, especially protein stabilization and protein-protein interaction ^4^. Therefore, SUMOylation plays indispensable roles in cellular signaling regulation. In cancer cells, it was discovered that SUMOylation enzymes were upregulated, indicating their beneficial contributions to cancer occurrence and development ^4^.

In previous studies, only one SUMO E1 (activating) enzyme was discovered, which was a heterodimer of Aos1 (or SAE1) and Uba2 (or SAE2). Inhibitors targeting SUMO E1 have great potential and has been actively pursued ^11–18^. Ginkgolic acid is the first chemical inhibitor of SUMOylation by inhibiting the formation of the SUMO E1-SUMO complex ^13^. ML-792 is an ATP mimics selectively inhibiting SUMO E1 ^17^. COH000 is the only covalent allosteric inhibitor (CAI) of SUMO E1. Compared to regular inhibitors, allosteric inhibitors have the advantage of low side-effect and high specificity. However, COH000 was discovered at a great cost from the experimental screening of over 290,000 compounds and extensive modifications ^11,19^.

This work is the first virtual screening of CAIs of SUMO E1. Compared to the discovery of COH000, we successfully identified two CAIs with new scaffolds and covalent warheads by experimentally testing only 13 compounds. Therefore, our work is highly efficient and cost-saving. Our work also largely expanded the chemical space for designing new allosteric SUMO E1 inhibitors. This is also the first time that SCARdock was used on the virtual screening of covalent inhibitors targeting PPI interface, demonstrating the great potential of this protocol in discovering diverse covalent ligands.

One limitation of this work is that we were not able to directly quantify the changes of Uba2-SUMO, Ubc9-SUMO, and MAT2A-SUMO. A major reason was that SUMOylation is a highly dynamic process and usually only a tiny fraction of a substrate protein is SUMOylated, which makes it very challenging to detect SUMOylated substrates in endogenous levels except RanGAP1^17^. We also failed to detect enough targets by immunoaffinity enrichment. Overexpressing the targets might be an option for further confirmation in the future. Another limitation is that we were not able to verify if these two new CAIs have off-target effects and could be further improved. Optimization of the structures of these two inhibitors would be necessary in future. Lastly, if the new scaffolds and covalent warheads are superior to COH000 was not thoroughly examined in this work.

Taken together, this work presented the first virtual high-throughput screening of SUMO E1 CAIs, successfully identified two new SUMO E1 CAIs with new scaffolds and warheads at low cost and validated the application of our covalent drug discovery protocol SCARdock on protein-protein interacting (PPI) interfaces for the first time. Our work also revealed a counter-intuitive crosstalk between the SUMOylation pathway and the polyamine metabolism pathway. This work would be of great value to the discovery of improved SUMO E1 inhibitors as well as covalent inhibitors targeting PPI interfaces.

## Material and Methods

### 1. Chemicals and reagents

The screening compounds were purchased from Mcule, Inc. (U.S.A). The compounds were validated by 1H NMR and/or LC/MS by the vendor and had at least 90% purity. The compounds were fully dissolved in DMSO before experimental assays. The rabbit polyclonal anti-Uba2 (Cat. NO. D151837) and anti-Ubc9 (Cat. NO. D262711) antibodies were from Sangon Biotech (Shanghai) Co., Ltd., China. The rabbit monoclonal anti-SUMO1 (Cat. NO. HY-P80351) was from MedChemExpress LLC (U.S.A.). The rabbit monoclonal anti-RanGAP1 (Cat. NO. R383099) and anti-MAT2A (Cat. NO. R389369) antibodies were from ZenBio, Inc. (U.S.A.). The rabbit polyclonal anti-AMD1 (AdoMetDC) (Cat. NO. DF12822) and anti-ODC (Cat. NO. DF6712) antibodies were from Affinity Biosciences (U.S.A.). The rabbit GAPDH Polyclonal antibody (Cat. NO. 10494-1-AP) was from Proteintech (U.S.A). The HRP-conjugated goat anti-rabbit IgG (Cat. NO. D110058) was from angon Biotech (Shanghai) Co., Ltd., China. The BeyoECL Plus supersensitive ECL chemiluminescence kit (Cat. NO. P0018AS) was from Beyotime Biotech (Shanghai) Co., Ltd., China. 3,5-Dimethoxy-4-hydroxycinnamic acid (sinapinic acid; SA) (Cat. NO. BD0554) was from Bide Pharmatech Co., Ltd., China, and MGBG (Mitoguazone) (Cat. NO. M193101) was from Shenzhen BSZH Design Co., Ltd., China. Putrescine (Cat. NO. V900377-25G), spermidine (Cat. NO. S2626-1G), spermine (Cat. NO. S2876-1G), 1,7-Diaminoheptane (DAH) (Cat. NO. D17408) were from Merck & Co. Inc., USA. The IBT-16plex kit (Cat. NO. K10000-16) was from Nanjing Apollomics Biotech Inc., China. COH000 (Cat. NO. C882038), trifluoroacetic acid (TFA) (Cat. NO. C11936727), dichloromethane (Cat. NO. D824411), and benzoyl chloride (Cat. NO. B802233) were from Macklin (Shanghai) Co., Ltd., China. S-Adenosyl-L-Methionine (AdoMet) (Cat. NO. S9990) was from Beijing Solarbio Science & technology (Beijing) Co., Ltd., China.

### 2. Cell culture

Human T47D breast cancer cells were purchased from the China Center for Type Culture Collection (Wuhan, China). The cells were authenticated with the short tandem repeat (STR) analysis by the vendor. The cells were kept in RPMI 1640 media supplemented with 10% fetal calf serum (FBS) (Upsilon Bio-tech Co., Ltd., USA) and 1% penicillin-streptomycin solution (P/S) (Gibco). The cells were cultured in a 37 °C incubator supplied with 5% CO_2_.

### 3. SCARdock screening

The SCARdock process was similar to what described previously ^27^. The complex structure of COH000 and the human SUMO E1 (PDB ID: 6CWY) was used as the target, with the covalent residue Cys30 being mutated to Gly in Pymol. The MCULE purchasable dataset (March 01, 2021) was filtered with the experimentally validated Cys-targeting warhead database we compiled and manually curated ^23^ before it was used in the SCARdock process using AutoDock Vina 1.1.2 ^46^. From 106,952 docked compounds, 2,026 hits were chosen for expert evaluation after being filtered with the following rules: (1) rmsd range relative to the first conformation: 3.0 Å; (2) the score of the first conformation: −8.0 kcal/mol; (3) percentage of the top 10 docked conformations fulfilling the rules (1) and (2): 0.75; (4) affinity density of the non-hydrogen atoms: −0.28 kcal/mol; (5) bonding atom distance: 2.0 Å; (6) the score of the conformation fulfilling the rule (5): −8.0 kcal/mol. After discarding hits with inappropriate warheads and considering warhead diversity, we selected 25 compounds for purchasing inquiry and 13 compounds were purchased for experimental validation.

### 4. Gene cloning

The coding DNA sequences of human Aos1 and SUMO1 were obtained by reverse transcription using a human RNA library stored in our laboratory. The coding DNA sequence of human Uba2 was synthesized by Wuhan Miaoling Biotechnology Co., Ltd., China. The coding sequences were digested (Aos1 and SUMO1: BamH I and Xho I; Uba2: Nco I and Xho I) and inserted into pET28a after being cloned by PCR with the following primers: Aos1-Forward, CGC GGA TCA CTC CGG GCG TGC TGC CG; Aos1-Reverse, CCG CTC GAG TCA CTT GGG GCC AAG GCA CTC CAC; Uba2-Forward, CAT GCC ATG GCA CTG TCG CGG GGG; Uba2-Reverse, CCG CTC GAG ATC TAA TGC TAT GAC ATC ATC AAG CTC T; SUMO1-Forward CGC GGA TCC ATG TCT GAC CAG GAG GCA AAA CCT TCA ACT GAG GA; SUMO1-Reverse, CCG CTC GAG CTA ACC CCC CGT TTG TTC CTG ATA AAC TTC AAT CAC. All sequences were verified by DNA sequencing.

### 5. Protein expression and purification

Aos1 and SUMO1 were constructed to be expressed with a N-terminal 6xHis tag, and Uba2 was constructed to be expressed with a C-terminal 6xHis tag. All proteins were expressed in *Escherichia coli* BL21 (DE3). Protein expression was induced with 1 mM isopropyl-β-D-1-thiogalactoside (IPTG) for 6 h at 25 °C when OD_600_ achieved 0.6-0.8. Cell pellets were harvested, resuspended, and lysed by sonication in the lysis buffer (50 mM NaH_2_PO_4_/ Na_2_HPO_4_, 350 mM NaCl, pH 8.0, 20 mM imidazole, 2 mM DTT, 0.01% Tween-20). To facilitate the complex formation of Aos1/Uba2 (SUMO E1), the cells expressing these two proteins were harvested and purified together as described in ^47^. The supernatant was separated by centrifugation and applied to Ni-NTA agarose (Qiagen) for purification. The target protein was eluted with the elution buffer (50 mM NaH_2_PO_4_/ Na_2_HPO_4_, 350 mM NaCl, pH 8.0, 250 mM imidazole, 2 mM DTT, 0.01% Tween-20). The eluted protein was further purified by gel filtration (Superdex 200; GE Healthcare) and eluted in the storage buffer (50 mM Tris-HCl, 50 mM NaCl, pH 8.0, 1 mM DTT). The proteins were flash frozen in liquid nitrogen and stored at −80 °C.

### 6. Mutagenesis of Uba2

The Cys30 of Uba2 was mutated into serine or alanine using the site-specific mutagenesis kit (TianGen Biochemical Technology Co., LTD., China). The mutagenesis primers were: Uba2-C30S-Forward, GCG GGC GGC ATC GGC TCC GAG CTC CTC AAG A; Uba2-C30S-Reverse, GAG CCG ATG CCG CCC GCC CCC ACC AC; Uba2-C30A-Forward, CGG CGC CGA GCT CCT CAA GAA TCT CGT GCT CAC CGG; Uba2-C30A-Reverse, CTT GAG GAG CTC GGC GCC GAT GCC GCC CGC CCC CAC. The mutation was verified with DNA sequencing. The mutant protein was expressed, purified, and stored as the wild-type Uba2.

### 7. SDS-PAGE gel analysis

The inhibition assay was performed by incubating 1 μM of the SUMO E1 (Aos1/Uba2 complex) with a compound at the indicated concentration in the reaction buffer (20 mM HEPES pH 7.5, 50 mM NaCl, 5 mM MgCl_2_) for the indicated time at 37 °C. ATP and SUMO1 were subsequently added to a final concentration of 1 mM and 2 μM respectively. Reactions continued at 37 °C for 1 h and terminated using the SDS-PAGE loading buffer. Samples were resolved on 12 % SDS-PAGE and visualized with Coomassie brilliant blue. The gel was photographed and the bands were quantitatively analyzed with ImageJ (NIH). All assays were performed at least three times.

### 8. Gel-based thermal shift assay

The protein sample was diluted to 1 mg/mL in the reaction buffer (20 mM HEPES pH 7.5, 50 mM NaCl, 5 mM MgCl_2_). Then, a 10 mM stock solution of the compound was added to a final concentration of 100 μM. The mixture was incubated at 37 °C for 1 h. Subsequently, the mixture was pipetted into PCR tubes (30 uL each) and heated at the indicated temperatures for 3 minutes in a PCR thermocycler. Then the heated samples were centrifuged at 12,000 rpm at 4 °C for 15 minutes, and 12 μL of the supernatant was analyzed with SDS-PAGE.

### 9. MALDI-TOF analysis

The compound was incubated with SUMO E1 (Aos1/Uba2) in the reaction buffer for 1 h at 37 °C before it was diluted with 0.1% TFA to 0.4-1.0 mg/mL. Then the sample was fully mixed with saturated SA matrix in a ratio of 1:1 (v/v). The evenly mixed solution was spotted on a MALDI anchor target plate (MTP 384 polished steel). The samples were subject to data acquisition on a high resolution UltrafleXtreme MALDI TOF mass spectrometer (Bruker Daltonics, Bremen, Germany) with Compass for flexSeries 2.0. Linear mode was used for ion sources (Ion Source 1: 19.5 kV, Ion Source 1: 16.6 kV) and Pulsed Ion Extraction (600 ns). Matrix suppression was set in Deflection mode to suppress matrix components up to 28000 Da. In the Sample Carrier, random walk constitutes a “Partial sample” where the number of shots at the raster spot was 10 and the limit diameter was 2000 μm. In detection, the mass range was 30000-210000 Da and detector gain was 18.5. Flexcontrol (v2.4) was used for data acquisition, and FlexAnalysis (2.4) was used for data processing.

### 10. Cell proliferation/cytotoxicity assay

T47D cells were resuspended and 100 µL of the suspension was inoculated in a 96-well plate at 0.6 × 10^5^ cells/mL per well. On the following day, cells were treated with 100 μL of medium containing the indicated compound. The final concentration of DMSO was 0.1% (v/v) for all samples. The medium was pipetted out after 24 h or 48 h of treatment, before 100 μL of medium containing 5% CCK-8 (water-soluble tetrazolium salt, WST-8) (Beyotime Biotech (Shanghai) Co., Ltd., China) was added. After 3h of incubation at 37 °C, the absorbance at 450 nm was measured with a microplate reader (BioTek Synergy H1).

### 11. Wound healing assay

T47D cells were resuspended and 100 µL of the suspension was inoculated in a 96-well plate at 2 × 10^5^ cells/mL per well. The next day, pipette tips were used to scratch the confluent monolayers in a straight line. The cells were then treated with 100 μL of medium containing the indicated compound. The final concentration of DMSO was 0.1% (v/v) for all samples. The cells were cultured in a cell incubator for 24 h before being photographed. The width of the wound area was measured with ImageJ and quantified using the following formula: mobility = [(scratch width at 0 h - scratch width at 24 h)/scratch width at 0 h] × 100%, relative mobility = (mobility of the treated sample/mobility of the control) × 100%. The width of the wound area was recorded as the average value of the two ends and the middle.

### 12. Western blotting

T47D cells were resuspended and 2 mL of the suspension was inoculated in a 6-well plate at 2 × 10^5^ cells/mL per well. On the next day, cells were treated with the indicated compound. The final concentration of DMSO was 0.1% (v/v) for all samples. After 48 h of treatment, cells were collected and lyzed with the RIPA lysis buffer. The protein concentration was quantified with BCA kit and analyzed by Western blotting. The dilution ratios of the antibodies were: Uba2, 1:1000; Ubc9, 1:1000; SUMO1, 1:1000; RanGAP1, 1:1000; MAT2A, 1:1000; AdoMetDC, 1:500; ODC, 1:500.

### 13. AdoMet quantification

T47D cells were resuspended and 2 mL of the suspension was inoculated in a 6-well plate at 2 × 10^5^ cells/mL per well. On the next day, cells were treated with the indicated compound. The final concentration of DMSO was 0.1% (v/v) for all samples. After 48 h of treatment, cells were collected and lyzed with three freeze/thaw cycles. The protein concentration was quantified with BCA kit. 100 µL of 40% (w/v) trichloroacetic acid was added into 50 µL of supernatant and the sample was placed on ice for 30 min to let the protein fully precipitate. Then 100 µL of ddH_2_O was added and the sample was centrifuged for 10 min at 4 °C, 15000 r/min. 500 µL of dichloromethane was added to the supernatant and the sample was vortexed for 2-3 min. The sample was centrifuged for 5 min at 12000 rpm, 4 °C, and the upper water phase was kept (extracted twice). The extracted sample was filtered through a 0.22 µm filter and analyzed by LC-MS.

### 14. Cellular polyamine quantification

The cells were treated as above. After the protein concentration was quantified with BCA kit, 20 µL of 50 µM DAH, 500 µL of 2 M NaOH, and 10 µL of benzoyl chloride were added into 200 µL supernatant. The mixture was vortexed for 30 seconds and then placed in a water bath at 42 °C for 20 min. Then, 2 mL of saturated NaCl aqueous solution was added to terminate the reaction, and 2 mL of dichloromethane was added for extraction for 1 min. After standing and layering, the lower organic phase was collected (extracted for three times). The collected phase was air dried and 1mL methanol was added to dissolve the sample, before the sample was filtered through a 0.22 µm filter and analyzed by LC-MS.

### 15. LC-MS

The cellular levels of AdoMet and polyamines were quantified on a QE Q Exactive UHMR Hybrid Quadrupole-Orbitrap MS System (Thermo Fisher). This system consisted of an ultra-high-performance liquid chromatography system coupled with a triple quadrupole mass spectrometer equipped with an ESI source. EclipsePlusC18 was used for chromatographic separation at 25 °C, and the flow rate was set at 0.2 mL/min. The sample injection volume was 10 µL. The mobile phase was water containing 0.1% formic acid (A) and methanol containing 0.1% formic acid (B).

For AdoMet, the elution program was set as: 0-1 min, 90% A; 1-3 min, 90-10% A; 3-6 min, 10% A; 6-6.01 min, 10-90% A; 6.01-8 min, 90% A. The MS parameters were: Ion transfer tube temperature, 350 °C; evaporation temperature, 300 °C; sheath gas pressure, 35 arb; auxiliary gas pressure, 10 arb; electric spray voltage, 3.5 kV; positive ion mode; scanning mode, targeted-SIM/dd-MS^2^; collision energy, NCE10; full scan resolution, 70000; automatic gain control (AGC), 5×10^4^; maximum injection time, 100 ms; dd-MS^2^ resolution, 35000; AGC, 2×10^5^; maximum injection time, 100 ms.

For polyamines, the elution program was set as: 0-1 min, 90% A; 1-5 min, 90-10% A; 5-8 min, 10% A; 8-8.01 min, 10-90% A; 8.01-10 min, 90% A. The MS parameters were: Ion source, HESI; positive ion mode; sheath gas flow rate, 40 arb; auxiliary gas presure, 10 arb; capillary temperature, 350 °C; electric spray voltage, 3.5 kV; scanning mode, full MS/dd-MS^2^; full scan resolution, 70000; automatic gain control (AGC), 5×10^4^; maximum injection time, 100 ms; dd-MS^2^ resolution, 35000; AGC, 2×10^5^; maximum injection time, 100 ms.

### 16. Quantitative 16-plex proteomic analysis

The proteomic analysis was performed as described in ^48^. In each well of a 100 mm cell culture dish, 5×10^6^ of T47D cells were seeded. On the following day, the cells were treated with the compounds for 48 h before the cells were collected and stored at −80 °C. Before proteomic analysis, the cells were resuspended in a lysis buffer containing 8 M urea, 20 mM Tris, pH 8.5, and protease inhibitor cocktail (Complete Mini, Roche). After sonication and centrifugation, the proteins in the supernatant were determined by the BCA assay. Then 100 µg of proteins was reduced with 10 mM dithiothreitol for 30 min at 37 °C and alkylated with 55 mM iodoacetamide for 45 min in dark. The treated proteins were diluted with 0.5 M TEAB (triethylammonium bicarbonate) and digested with trypsin (Prometa) at a ratio of 30:1 (protein/enzyme, w/w) for 12 h at 37 °C. The digested sample was desalted on a MonoSpin C18 (GL Sciences) column and lyophilized through a ScanSpeed 40 ScanVac Vacuum Concentrator (ALEX RED). The digested peptides (50 µg) were then labeled with IBT-16plex according to the instruction. Then, the labeled peptides were pooled, desalted, and fractionated by HPLC (1260 Infinity, Agilent) with a Gemini C18 column (5 µm, 4.6 × 250 mm). Briefly, the gradient parameters were set as 5% acetonitrile (pH 9.8, Solvent A) and 95% acetonitrile (pH 9.8, Solvent B): 0-10 min, 5% B; 10-40 min, 5-35% B; 40-44 min, 35-95% B; 44-45 min, 95-5% B; 45-60 min, 5% B. Peptides were fractionated into 60 fractions in 60 min and combined into 12 fractions, which were then subjected to vacuum-dried. The labeled peptides were detected by an Orbitrap Eclipse mass spectrometer (Thermo Fisher) coupled with a nano reversed phase (RP) column (5 μm, Hypersil C18, 75 μm × 150 mm), with setting in positive ion mode and data-dependent manner with full MS scan from 400 to 1600 m/z, resolution at 120000, MS/MS scan with minimum signal threshold 20000, and isolation width at 2 Da except with an MS/MS resolution of 35000. The data of each sample were collected from two technical replicates.

### 17. Gene expression analysis of MAT2A

DepMap (https://depmap.org/portal/) was used to assess the importance of MAT2A in different cancer types. The MAT2A CRISPR Dependence Scores (CRISPR (DepMap Public 24Q2+Score, Chronos)) were retrieved from the DepMap ^49^ portal and plotted. The Kaplan-Meier plotter server (http://kmplot.com/analysis/) was used to analyze the expression of MAT2A in Luminal A (2013 StGallen criteria) breast cancer patients. The median value was used in the stratification of patients into groups with high and low MAT2A expression levels. A Kaplan-Meier survival curve was subsequently generated to compare the recurrence-free survival probabilities between these two groups. The statistical significance was assessed using the Log-rank test.

### 18. SUMOylation prediction

The SUMOylation sites of the proteins were predicted using the GPS-SUMO 2.0 server (https://sumo.biocuckoo.cn/) ^45^. The thresholds for both SUMOylation and SUMO interaction were set as high.

### 19. Statistical analysis

GraphPad Prism 8.2.1 was used for the statistical analyses unless indicated otherwise. The proteomic data were analyzed in R. Data were shown as means ± standard deviation (SD). Significance analyses were carried out using Student’s t-test. The threshold for statistical significance was p ≤ 0.05.

## Author contributions

Conceptualization, S.L.; methodology, S.Z., Y.T.J., X.Y.S., and S.L.; software, S.L. and X.Y.S.; validation, S.Z.; formal analysis, S.Z. and S.L.; investigation, Z.S., S.L., Y.T.J., X.Y.S., and Z.Y.W.; resources, S.L.; writing-original draft preparation, S.Z. and S.L.; writing-review and editing, S.L.; visualization, S.Z., X.Y.S., and S.L.; supervision, S.L., Y.B.A., and J.P.W.; project administration, S.L.; funding acquisition, S.L.. All authors have read and agreed to the published version of the manuscript.

## Additional information

### Competing financial interests

The inhibitors discovered in this work have been submitted for patent application in China.

## Acknowledgements

We would like to thank the support from the other members of our lab. This research was funded by the National Natural Science Foundation of China grant number [31971150], the Department of Science and Technology of the Hubei Provincial People’s Government grant numbers [2024AFA014, 2019CFA069], and the Opening Fund of Collaborative Innovation Center for Industrial Fermentation (Ministry of Education & Hubei Province) grant number [2023SL01].

## Data availability

The Supporting Information is available free of charge at the web site.

