## Supplementary Information for "SUMO E1 covalent allosteric inhibitors modulate polyamine synthesis via the MAT2A-AdoMetDC axis"

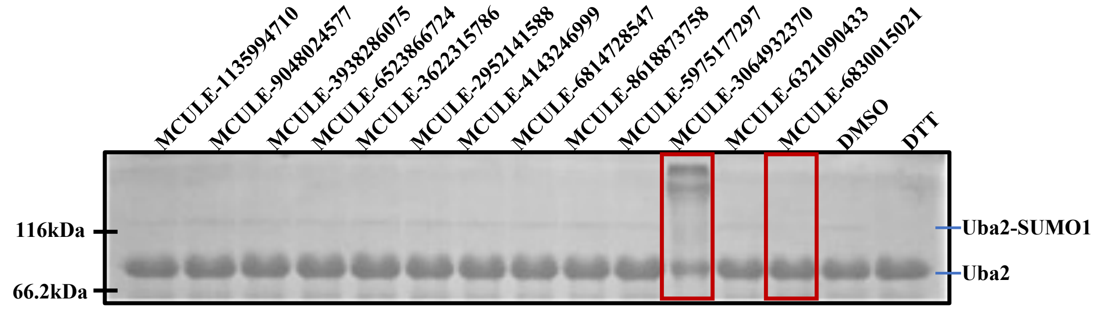


**Figure S1**. MCULE-3064932370 and MCULE-6830015021 inhibited the formation of the SUMOylation of Uba2. Before the addition of SUMO1, the compounds were pre-incubated with Aos1/Uba2 for 60 minutes at various concentrations according to their solubility. MCULE-3064932370 and MCULE-6830015021 were 100 µM, MCULE-6321090433 was 50 µM, and the others were 500 µM. The reductant DTT (dithiothreitol) inhibits SUMOylation and was used as a control. All samples had 1% (v/v) DMSO. The high molecular-weight bands in the MCULE-3064932370 lane were uncharacterized.

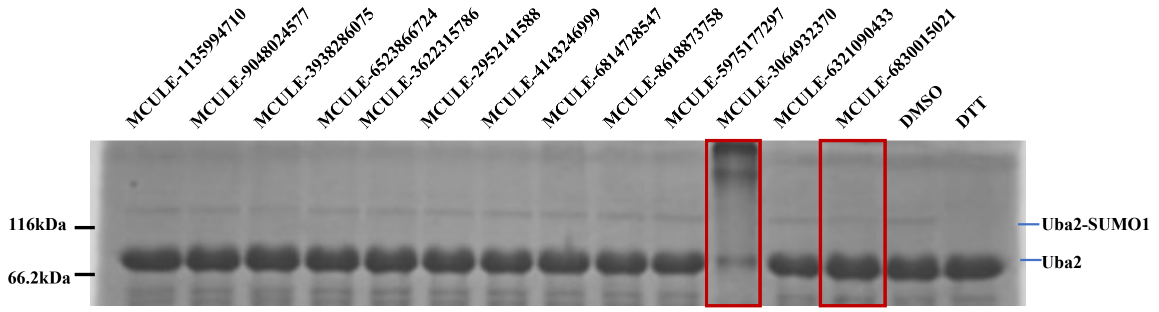


**Figure S2**. MCULE-3064932370 and MCULE-6830015021 inhibited the formation of the SUMOylation of Uba2. Before the addition of SUMO1, the compounds were pre-incubated with Aos1/Uba2 for 90 minutes at various concentrations according to their solubility. MCULE-3064932370 and MCULE-6830015021 were 100 µM, MCULE-6321090433 was 50 µM, and the others were 500 µM. The reductant DTT (dithiothreitol) inhibits SUMOylation and was used as a control. All samples had 1% (v/v) DMSO. The high molecular-weight bands in the MCULE-3064932370 lane were uncharacterized.

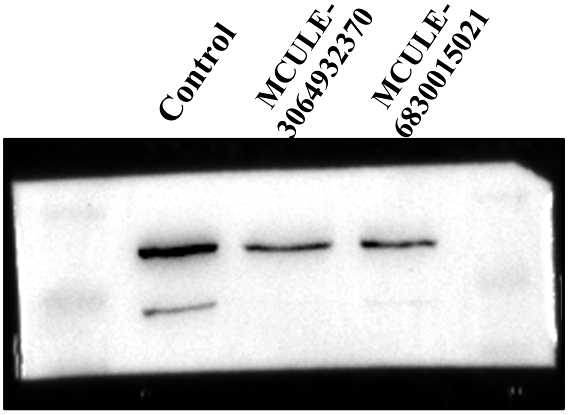


**Figure S3**. The full gel of the Western blotting data of RanGAP1 shown in Figure 4a.

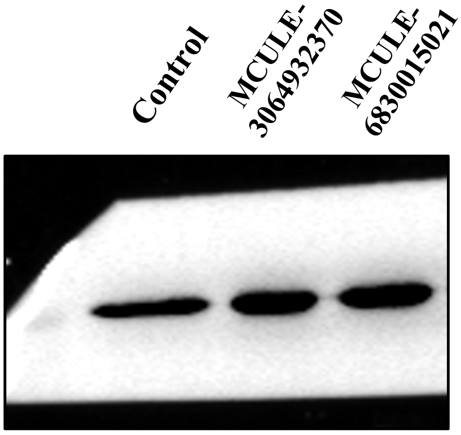


**Figure S4**. The full gel of the Western blotting data of GAPDH shown in Figure 4a.

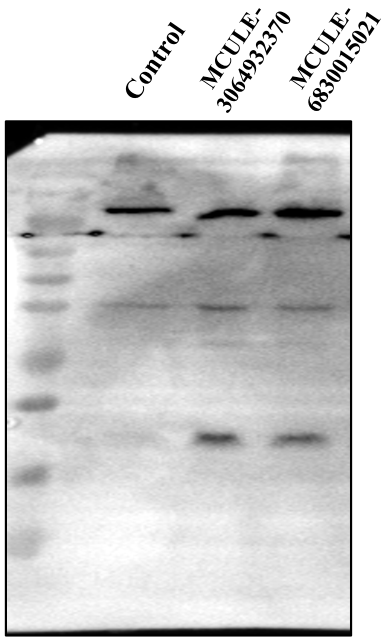


**Figure S5**. The full gel of the Western blotting data of SUMO1 shown in Figure 4a.

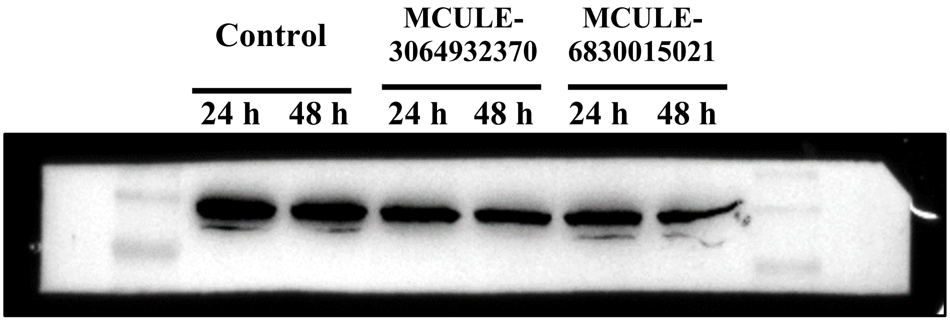


**Figure S6**. The full gel of the Western blotting data of Uba2 shown in Figure 4b.

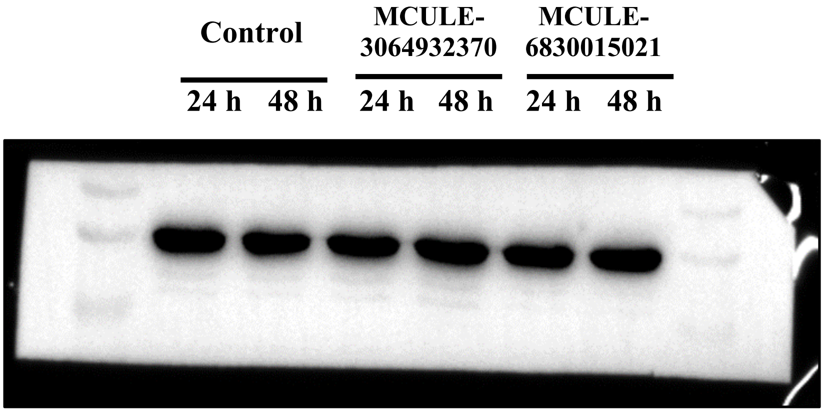


**Figure S7**. The full gel of the Western blotting data of GAPDH shown in Figure 4b.

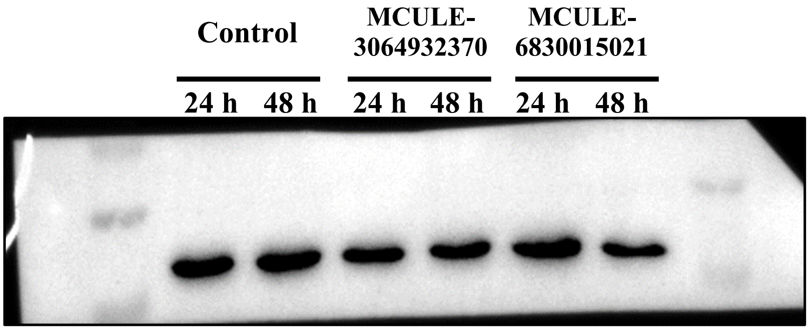


**Figure S8**. The full gel of the Western blotting data of Ubc9 shown in Figure 4b.

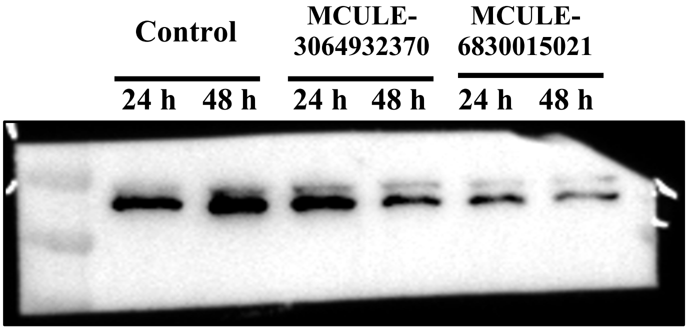


**Figure S9**. The full gel of the Western blotting data of MAT2A shown in Figure 5b.

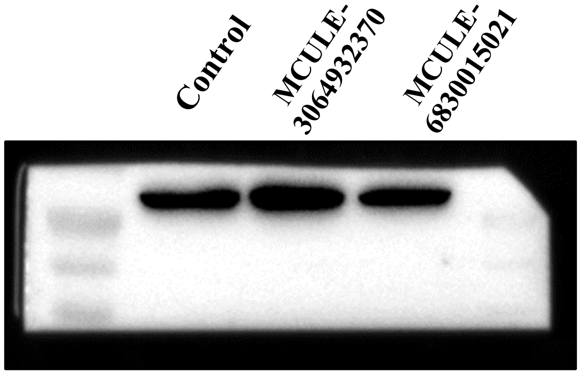


**Figure S10**. The full gel of the Western blotting data of ODC shown in Figure 5e.

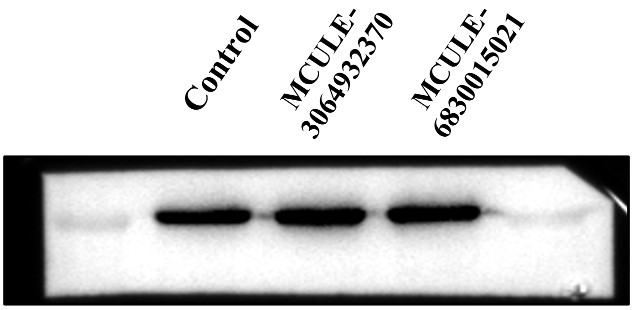


**Figure S11**. The full gel of the Western blotting data of GAPDH shown in Figure 5e.

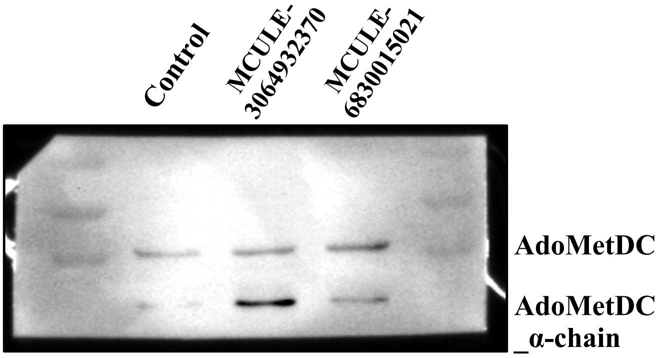


**Figure S12**. The full gel of the Western blotting data of AdoMetDC shown in Figure 5e.

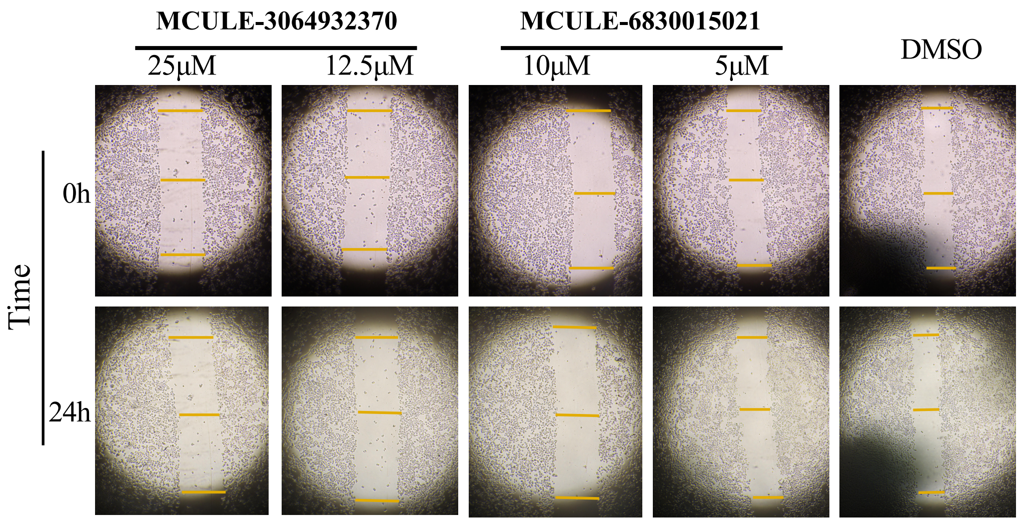


**Figure S13**. Representative cell migration images for the statistacal data in Figure 6d.
